# First natural crossover recombination of intact ORFs between two distinct species of the family *Closteroviridae*

**DOI:** 10.1101/325142

**Authors:** Leticia Ruiz, Almudena Simón, Carmen García, Leonardo Velasco, Dirk Janssen

## Abstract

*Lettuce chlorosis virus*-SP (LCV-SP) (family *Closteroviridae*, genus *Crinivirus*), is a new strain of LCV which is able to infect green bean plants and incapable of infecting lettuce crops. In the present study, high throughput and Sanger sequencing of RNA was used to obtain the LCV-SP full-length sequence. The LCV-SP genome comprises 8825 nt and 8672 nt equivalent with RNA1 and RNA2 respectively. RNA1 of LCV-SP contains four ORFs, the proteins encoded by the ORF1a and ORF1b are closely related to LCV RNA1 from California (FJ380118) whereas the 3´ end encodes proteins which share high amino acid sequence identity with RNA1 of BnYDV (EU191904). The genomic sequence of RNA2 consists of 8 ORFs, instead of 10 ORFs contained in LCV-California isolate. The distribution of vsiRNA (virus-derived small interfering RNA) along the LCV-SP genome suggested the presence of subgenomic RNAs corresponding with HSP70, P6.4 and P60. Results of the analysis using RDP4 and Simplot programs are the proof of the evidence that LCV-SP is the first recombinant of the family *Closteroviridae* by crossover recombination of intact ORFs, being the LCV RNA1 (FJ380118) and BnYDV RNA1 (EU191904) the origin of the new LCV strain. Genetic diversity values of virus isolates in the recombinant region obtained after sampling LCV-SP infected green bean between 2011 and 2017 might suggest that the recombinant virus event occurred in the area before this period. The presence of LCV-SP shows the role of recombination as a driving force of evolution within the genus *Crinivirus*, a globally distributed, emergent genus.

## Introduction

*Lettuce chlorosis virus* (LCV) belongs to the genus *Crinivirus*, family *Closteroviridae*. Viruses included in this family are the largest among the known plant viruses and present ssRNA, positive-sense genome [1]. Family *Closteroviridae* has been classified in three genera based on vector transmission and phylogenetic relationships: *Closterovirus, Ampelovirus*, and *Crinivirus*. Recently, a new genus named *Velarivirus* has been proposed [2]. All members of genus *Crinivirus* include segmented genomes, are whitefly-transmitted and limited to the phloem [3].

Many members of genus *Crinivirus* are considered emerging epidemics worldwide [4]. LCV was described in the 1990´s in California, transmitted by the whitefly *Bemisia tabaci* and affecting lettuce and sugar beet crops [5]. The complete sequence and the genomic organization of LCV were described in 2009 by [6] and comprises a bipartite genome formed by RNA1 and RNA2. LCV RNA1 contains the replication module (ORFs 1a and 1b), which encodes conserved domains of a papain-like leader proteinase (P-PRo), a methyltransferase (MTR), helicase (HEL) and RNA-dependent RNA polymerase (RdRp), ORF2 encodes a putative 8-kDa protein (P8), and ORF3 encodes a 22.9-kDa protein (P23) recently described as viral suppressor of RNA silencing [7]. LCV RNA2 contains the hallmark *Closterovirus* gene array coding for a heat shock protein homolog (Hsp70h, ORF3), a 50-60 kDa protein (ORF5), the major (CP, ORF7) and the minor coat protein (CPm, ORF 8) [1,8]. Furthermore, P5.6, P6, P6.4, P9, P27, and P4.8 are encoded by ORF1, ORF2, ORF4, ORF6, ORF9 and ORF10, respectively; the functions of these putative proteins remain unknown [6].

LCV share high homology with *Cucurbit chlorotic yellow virus* (CCYV) and with *Bean yellow disorder virus* (BnYDV), [9,6], this last one, was recognized as the first *Crinivirus* that infects Leguminosae family crops [10]. LCV was considered as an endemic infection which had not spread to areas outside Southwestern USA [4]. In 2014 however, symptoms similar to BnYDV in green bean crops were attributed to a new strain of LCV, LCV-SP, which was unable to infect lettuce plants and capable of affecting green bean crops [11].

Viral RNA-RNA recombination and/or reassortment of genomic segments (pseudo-recombination), between virus strains or virus species, increase the genetic variability and is a powerful tool in virus evolution, responsible for the emergence of new viral strains or species [12]. RNA recombination can also be used to repair viral genomes which will contribute to the fitness of viral populations. The low frequency of its detection is probably the result of selection pressure that removes the majority of recombinants [13]. The first evidence of genetic recombination was described in *Bromo mosaic virus* [14] and it seems to be specially frequent among species belonging to the family *Potyviridae* where crossovers have been described at ORFs and 3´UTR sequences [15–17,13].

Within family *Closteroviridae* homologous recombination has been described at CP and P20 in the *Closterovirus* member *Citrus tristeza virus* (CTV) [18,19] and at CP of the *Ampeloviruses* Grapevine leafroll-associated virus 3 (GLRaV-3) and GLRaV-11 [19,20]. Although *Closterovirus* members share a core of conserved genes, they present high variability in the set of ORFs located in the 3´end of RNA or in the 3´end of RNA 1 if the genome is bipartite, as is the case of genus *Crinivirus* [1,21]. The presence of mechanisms capable of increasing genetic variability such as genome recombination may have important evolutionary implications because some of these proteins have been characterized as RNA-suppressing proteins (RSS)[22–25,7].

In this study we describe a recombinant *Crinivirus*, LCV-SP, originating from a crossover of intact ORFs and analyse a population of isolates collected in the area. Sequencing was achieved by combining primer walking strategy and high-throughput sequencing. The implications in emergence of *Crinivirus* epidemiology are discussed.

## Material and methods

### Virus isolate

Between 2011 and 2017 samples of green bean leaves showing symptoms that could be attributed to LCV-SP (interveinal mottling and yellowing in lower and middle leaves [11]) were collected from different geographical locations from Granada, Almeria and Málaga provinces in Southeastern Spain. The plants were analysed by RT-PCR to ensure the presence of LCV-SP [11] and stored at −80°C for subsequent molecular analysis. One of these isolates collected in 2013 and annotated as “Almeria” was used for complete genome determination.

### Whole-genome sequencing of LCV-SP and genome analysis

Total RNA from LCV-SP isolate “Almeria” was extracted with Trizol Reagent (Invitrogen) according to the manufacturer’s instructions. Partial length viral sequence of RNA1 and RNA2 of LCV genome was determined by RT-PCR using a primer walking strategy with a set of degenerated primers based on LCV (FJ380118, FJ380119) and BnYDV (NC010560, NC010561) sequences (data not shown). Partial sequences of LCV-SP RNA1 and RNA2 previously described [11] were also used to design specific primers and flank the gaps. The sequences were assembled using the software GENEIOUS (v. 7.1.9, Biomatters, New Zealand).

### Small RNA purification, dsRNA extraction and deep sequencing

Small RNA purification from the Almeria LCV-SP isolate was obtained using the miRCURY RNA isolation kit (Exiqon, Denmark), according to the manufacturer’s instructions. Small RNA library was prepared using the Illumina TruSeq Small RNA Kit (Illumina, San Diego, California, USA). The library was sequenced on the Illumina deep sequencing platform using the services provided by CRG (Barcelona, Spain). Sequencing adapters and reads shorter than 18 nucleotides (nt) were removed using GENEIOUS and reads ranging from 18 to 24 nt were selected. These reads were assembled into contigs larger than 33 nt using the *de novo* assembly function of VELVET v. 1.2.08 implemented in the SCBI Picasso server (http://www.scbi.uma.es) with a k-mer = 18. Partial RNA1 and RNA2 LCV-SP sequence obtained for Sanger sequencing was used as a reference for contigs sequences. RNA1 and RNA2 consensus draft sequences obtained were ensured through RT-PCR amplification using specific primer pair based on the assembled genomes (not shown). The 5´/3´ Smart™ RACE cDNA Amplification Kit (Clontech, California, USA) was used to complete the 5´ and 3´ends. PCR fragments were cloned in pGemT vector and at least two cDNA clones were sequenced in both directions. To obtain the profile distribution of LCV-SP virus-derived small interfering RNAs (vsiRNAs), the 21 and 22 nt reads of the small RNA library were aligned to the assembled LCV-SP genome using MISIS-2 [26].

### Double stranded RNA purification

Deep sequence from LCV-SP dsRNA was used to confirm the LCV-SP RNA1 and RNA2 genomic sequences. For that, double-stranded RNA was purified from another LCV-SP isolate collected in 2014 [27] followed by S1 nuclease (Promega, Madison, Wisconsin, USA) and DNAase I (Invitrogen, Carlsbad, California, USA) treatments to discard the contamination with genomic DNA and ssRNAs. A library was constructed and sequenced using the Illumina deep sequencing platform with the services provided by CRG (Barcelona, Spain). Contigs where assembled after trimming the low-quality reads by using VELVET (k-mer = 31). Complete sequence of LCV-SP obtained as above is described was used as reference for contigs aligning in GENEIOUS v. 7.1.9.

The contigs obtained from both massive deep sequencing were conducted to BLAST analysis against the GenBank database in NCBI (http://www.ncbi.nlm.nih.gov) using GENEIOUS. The e-value cut-off was set to 10^−5^ so high confidence matches could be reported. The numbers of reads aligning to LCV or BnYDV genomes were annotated to confirm the presence of sequences belonging to both virus species.

### Phylogenetic analysis

For phylogenetic analysis, the complete genome sequences from 9 members of genus *Crinivirus*, retrieved from the GenBank database and the complete sequence of LCV-SP were compared. The pairwise percentage of nucleotide sequence identity was performed using the Clustal W algorithm present in MEGA v 7.01 program [28]. Appropriate nucleotide substitution models for each genomic region were determined using jModelTest 2.1 [29] and the best model proposed by the Akaike information criterion (AIC) applied in each cases (GTR+I+G for RNA1 and HSP70h; JC for RNA2, GTR+G for CP, CPm and RdRp). Bayesian consensus phylogenetic trees were inferred using Bayesian inference and Markov chain Monte Carlo (MCMC) simulation implemented in MrBayes plugin. MCMC analyses were run with a chain length of 10^7^ generations, sampling every 1000 trees and with a burn-in of 25% and chain heated to 0.2.

## Recombination analysis

Recombination Detection Program 4 (RDP4) was used to detect potential recombination events in LCV-SP, likely parental sequences, and localization of recombination breakpoints [30]. The program was executed with default parameter settings which include the RDP, GENECONV, Chimaera, MaxChi, BOOTSCAN, and SISCAN methods. For recombination analysis, sequences of RNA1 and RNA2 of LCV, BnYDV, LCV-SP and CCYV were aligned in MEGA v.7.0.1 software [28] and exported to the RDP4 program for recombination analysis. The program was performed using the default settings and a Bonferroni corrected *p*-value cut-off (α=0.05). Only recombination events detected by four or more methods were considered as significant. The results obtained by RDP were confirmed using a boot scanning method in the SimPlot program v.3.5.1 [31].

### Genetic diversity in the recombinant genomic region

The crossover recombination event was confirmed in several isolates by RT-PCR. Eleven isolates obtained in the surveys from 2011 to 2017 were analysed by RT-PCR after RNA extraction using the primers LCVSP-44F 5´GCATTCAAGAAATTGTGGGATG 3’ (nt 7517-7538) and LCVSP-50R 5´ATATTAATGTAATTCTACGGTC 3´ (nt 8738-8759) which amplified a region of the LCV-SP genome encompassing 1242 bp corresponding to the entire region of the p26 and p6 proteins. Reverse transcription of the isolates was carried out with a reverse primer using Superscript II (Invitrogen) at 42ºC for 60 min. Subsequently, DNA fragments were amplified by PCR with Expand High Fidelity PCR System (Roche, Basel, Switzerland) under the following conditions: 95ºC for 3 min, 35 cycles of denaturation for 20s at 95°C, annealing for 30s at 55ºC and extension for 40s at 72ºC followed by one final extension cycle for 5 min at 72ºC. PCR products were cloned into pGemT-Easy vector (Promega Corporation, Madison, USA). The clones were bidirectionally sequenced. Nucleotide alignments were performed using Clustal W in MEGA v. 7.01 [28] software with default settings. The aligned sequences were used to calculate the genetic distances for synonymous (dS) and non-synonymous substitution (dNS) as described by [32,33] (PBL method).

### Molecular differentiation between yellowing induced by BnYDV or LCV recombinant virus (LCV-SP)

In order to determine analytically the presence of the recombinant *Crinivirus* LCV-SP, a RT-PCR-restriction fragment length polymorphism (RFLP) was performed. The set of primers LC-Bn56F 5´GATTTTGGATTTGAAGC 3´ and LC-Bn51R 5´ACAACAGATCAAAATCCACAATG 3´ which amplified fragments of LCV-SP RNA1 and BnYDV RNA1 genome of 739 and 769 bp, corresponding to nucleotides 7190-7928 and 7294-8062 respectively were used for a RT-PCR reaction as described above. After that, the PCR products were digested with *Kpn*I (Promega) following the manufacturer’s instructions. Band patterns were analysed on a 2% agarose gel.

## Results

### Genome sequencing of LCV-SP

Walking strategy and Sanger sequencing allowed us to obtain 7109 bp and 7821 bp corresponding to 80% and 90% of complete genome of RNA1 and RNA2 of LCV-SP. The LCV-SP genome was completed by deep sequencing from small RNAs and RACE strategy. The genome was confirmed through RT-PCR amplification with specific primers (S1 Table) and consisting of 8825 nt and 8672 nt equivalent with RNA1 and RNA2 of GenBank accession numbers MG489894 and MG489895, respectively. Genetic organization of LCV-SP genome is shown in Fig 1. RNA1 of LCV-SP contains four ORFs: ORF1a (nt 73-7561) and ORF1b (nt 6044-7561) contains the replication module and the P-PRO, MTR and RdRp motifs. ORF2 (nt 7637-8338) encodes a predicted protein of 26-kDa (P26) and it is followed by another putative protein of 6-kDa (P6) (ORF3, nt 8339-8512). The proteins encoded by ORF1a and ORF1b share a 93.6 % and 99.6% respectively of amino acid sequence identity with LCV RNA1 from California (FJ380118). Whereas the 5´ from LCV-SP RNA1 are closely related to the LCV RNA1 genome previously described; the 3´ end encodes proteins corresponding to the RNA1 of BnYDV (EU191904). The P26 (ORF2) and P6 (ORF3) share a 99.5% and 100% amino acid sequence identity with those present in BnYDV. Both proteins currently have an unknown function; P26 has no homologues in the GenBank database and P6 display only 30% of amino acid sequence identity with a P6a protein of a new unclassified *Crinivirus* named *Tetterwort vein cholorosis virus* (ALE18214). The genomic sequence of RNA2 consists of 8 ORFs, instead of the 10 ORFs contained in LCV-California isolate [6].

The genomic sequence comprises the hallmark gene array of the family *Closteroviridae* coding for a HSP70h (ORF5, nt 1817-3483), and three diverged copies of the capsid protein: the ortholog of the coat protein homolog (ORF4, nt 3642-5195), the CP (ORF6, nt 5537-6289) and the CPm (ORF7, nt 6289-7713) which exhibit more than 95% amino acid sequence identity with RNA2 of LCV FJ380119. RNA2 of LCV-SP encodes also three putative small proteins: P5.6 (ORF1, nt 254-421), P6.4 (ORF3, nt 3484-3648) and P9 (ORF5, nt 5177-5416), which show 92%, 98% and 96% nucleic acid identity with those in LCV FJ380119. Two putative small proteins in the RNA2 of LCV-Californian isolate (FJ380119), P6 in the 5´ end, and P4.8 in the 3´end, which have unknown function are absent in LCV-SP. The last ORF of RNA2 (ORF9) encodes a putative P27, which shows a high amino acid identity (98%) with its counterpart protein in LCV FJ380119.

**Fig 1.**
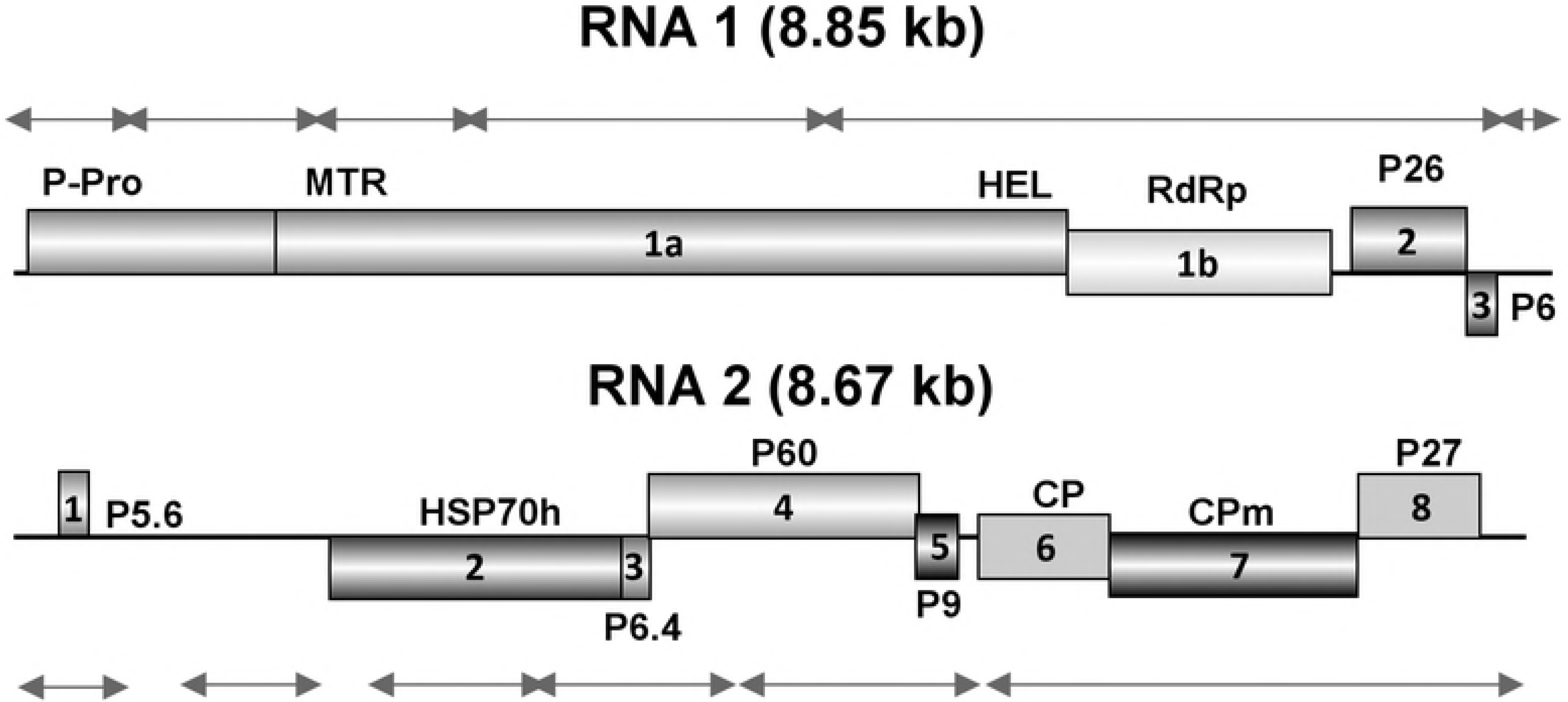
Diagram of genome organization of LCV-SP genome. Boxes represent ORFs; putative protein products are also indicated. Solid lines represent the location of contigs assembled from virus-derived small RNAs of LCV-SP.

### 5´and 3´ non coding region

The putative 5’UTR of LCV-SP RNA1 and RNA2 are 72 and 269 nt in length, and share 99% and 96% of nucleotide identity with LCV FJ380118/FJ380119 respectively. The first 5 nucleotides, GAAAT, are identical in both RNAs. This feature has been described in the Californian LCV isolate and in other members of genus *Crinivirus* [6]. Table 1 shows the percentage of nucleotide identity among 3´UTR regions corresponding with LCV-SP (RNA1 and 2), BnYDV (RNA1 and 2) and the Californian isolate of LCV (RNA 1 and 2). The 3´UTR of RNAs 1 and 2 are 313 and 256 nt in length respectively and shared high nucleotide sequence similarity (72%). Also, 3´UTR of RNA1 displays almost identical nucleotide sequence (99%) as its equivalent in BnYDV. This sequence is next to the putative proteins P26 and P6 corresponding to the 3´end of RNA1 of BnYDV and LCV-SP. Additionally, 3´UTR of RNA2 of LCV-SP show the highest homology (89%) with the 3´UTR of RNA1 of the Çalifornian isolate of LCV FJ380118.

**Table 1.** Percentage of nucleotide identity between 3´UTRs region of LCV-SP, LCV and BnYDV.

|  | LCV-SP RNA1 | BnYDV RNA1 | LCV RNA1 | LCV-SP RNA2 | LCV RNA2 | BnYDV RNA2 |
| --- | --- | --- | --- | --- | --- | --- |
| LCV-SP RNA1 |  | 99.7 | 70.1 | 71.4 | 72.2 | 56.2 |
| BnYDV RNA1 | 99.7 |  | 69.8 | 71.4 | 71.4 | 57.0 |
| LCV RNA1 | 70.1 | 69.8 |  | 89.5 | 79.8 | 57.8 |
| LCV-SP RNA2 | 71.4 | 71.4 | 89.5 |  | 79.6 | 61.3 |
| LCV RNA2 | 72.2 | 71.4 | 79.8 | 79.6 |  | 52.8 |
| BnYDV RNA2 | 56.2 | 57.0 | 57.8 | 61.3 | 52.8 |  |

LCV RNA1: FJ380118; LCV RNA2: FJ380119; BnYDV RNA1: EU191904; BnYDV RNA2: EU191905; LCV-SP-RNA1: MG489894; LCV-SP RNA2: MG489895

### Libraries of small RNAs and dsRNAs

After removal of adapters and reads shorter than 18, deep sequencing from small RNAs library produced a set of 44,578 million reads, which were subjected to the novo assembled, generating 19,739 contigs. A total of 113 and 125 sequences of these contigs assembled with LCV-SP RNA1 and RNA2, respectively. Fig 1 shows the location of the contigs derived from small RNA reads and assembled to the LCV-SP RNA1 and RNA2 genome components. The recombinant genome of LCV-SP was confirmed also by deep sequencing library obtained from dsRNA extracted from a LCV-SP green bean-infected plant that resulted in 31,353,455 reads. These sequences were assembled resulting in 3642 contigs. These contigs covered 90% of the LCV-SP RNA1 genome and the consensus sequence result was 100% identical. Contigs assembled with LCV-SP RNA2 covered 94% of LCV-SP RNA2 genome with 99.7% coincidence.

The result of BLASTN is shown in S2 Table. Only sequences matching the RNA1 and RNA2 of different LCV isolates and the 3´end of the RNA1 of BnYDV (EU191904), corresponding to P26 and the 3´UTR, were identified. All the sequences identified in BLASTN as part of the genome of RNA1 of LCV showed high nucleotide identity (more than 80%) with ORFs 1a and 1b. No sequences corresponding to the 3´end of RNA1 of LCV were identified, confirming the presence of the first *Crinivirus* arising by crossover recombination of entire ORFs and discarding the presence of a native LCV in the plants. The profiles of vsiRNAs aligned to LCV-SP RNA1 and 2 were analysed. The 21-nt vsiRNA class population was most prevalent, followed by the 22-nt (10.1 and 2.6 million respectively.) From this 21-nt population, 58 675 and 80 904 reads matched to LCV-SP RNA1 and 2 genome. In the case of 22-nt class population, 53.932 and 68.146 matched with LCV-SP RNA1 and 2. The analysis of 21 and 22-nt (not shown) vsiRNA strand polarity showed similar pictures, 53% of 21 and 22-nt vsiRNA class populations aligned to LCV-SP RNA1 were negative, whereas more than 50% of 21-nt and 22-nt vsiRNA class populations aligning to LCV-SP RNA2 were positive (53% and 51%, respectively). The profile distribution of both vsiRNAs populations aligning to the LCV-SP RNA1 and RNA2 genome components were also very similar. The distribution of 21-nt vsiRNAs along LCV-SP RNA1 and 2 genome is shown in the Fig 2. Several vsiRNA-generating regions (named as hot spots) were identified in both genomic RNAs of LCV-SP. Hot spots in RNA1 were located between MTR and HEL domains. The distribution of vsiRNA hotspots in the RNA2 genome corresponding with ORF 2 (HSP70), 3 (P6.4) and 4 (P60) are correlated with the position of subgenomic RNAs in LCV described by [6]. A hotspot located in the position 1361 of RNA2 does not correspond with any putative coding region.

**Fig 2.**
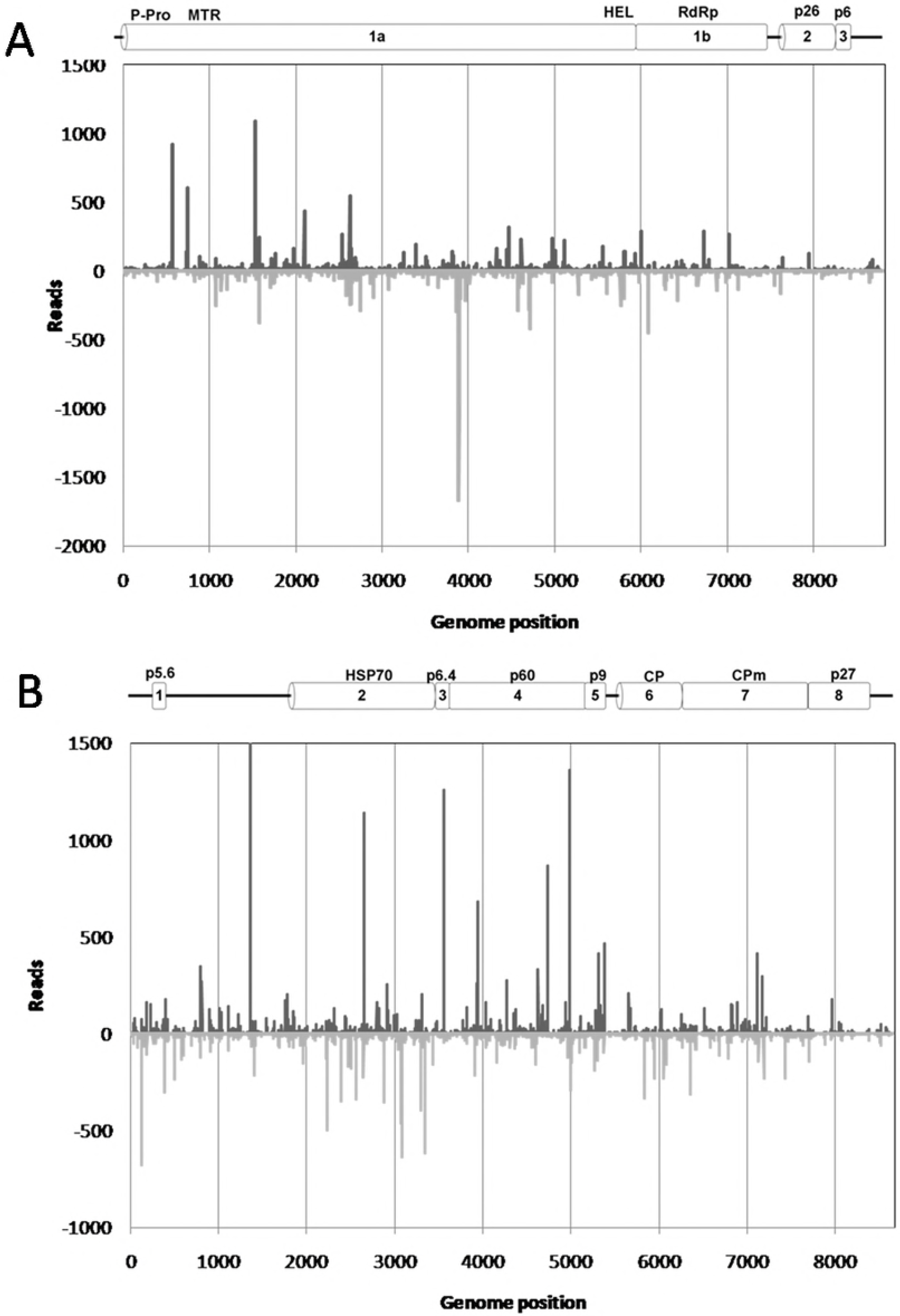
Profiles distribution of 21-nt vsiRNAs along LCV-SP RNA1 (A) and RNA2 (B). Sense and antisense of 21-nt vsiRNAs orientations are above and below the axis respectively. Schema of LCV-SP RNA1 and RNA2, genome is represented as reference.

Investigating the 5´-and 3´-terminal nucleotides of the vsiRNAs provides insight into the specific silencing machinery active in the plant. In S1 Fig we represent the percentages of nucleotides at each terminal position for the 21 to 24-nt vsiRNA populations matching LCV-SP RNA1 and RNA2. For both RNAs, there is a bias towards A/U in the 5´ and 3´ nucleotide positions in the 21 to 24-nt classes of vsiRNAs.

### Phylogenetic analysis

The phylogenetic trees generated with the complete sequence of RNA1 and RNA2 of LCV-SP and other members of genus *Crinivirus* show two well differentiated lineages; one comprising LCV-SP, LCV, CCYV, BnYDV and CYSDV and the other is formed by BPYV, ToCV, PVYV and LIYV (Fig 3) Then, two clades were generated where the divergence is clear.

**Fig 3.**
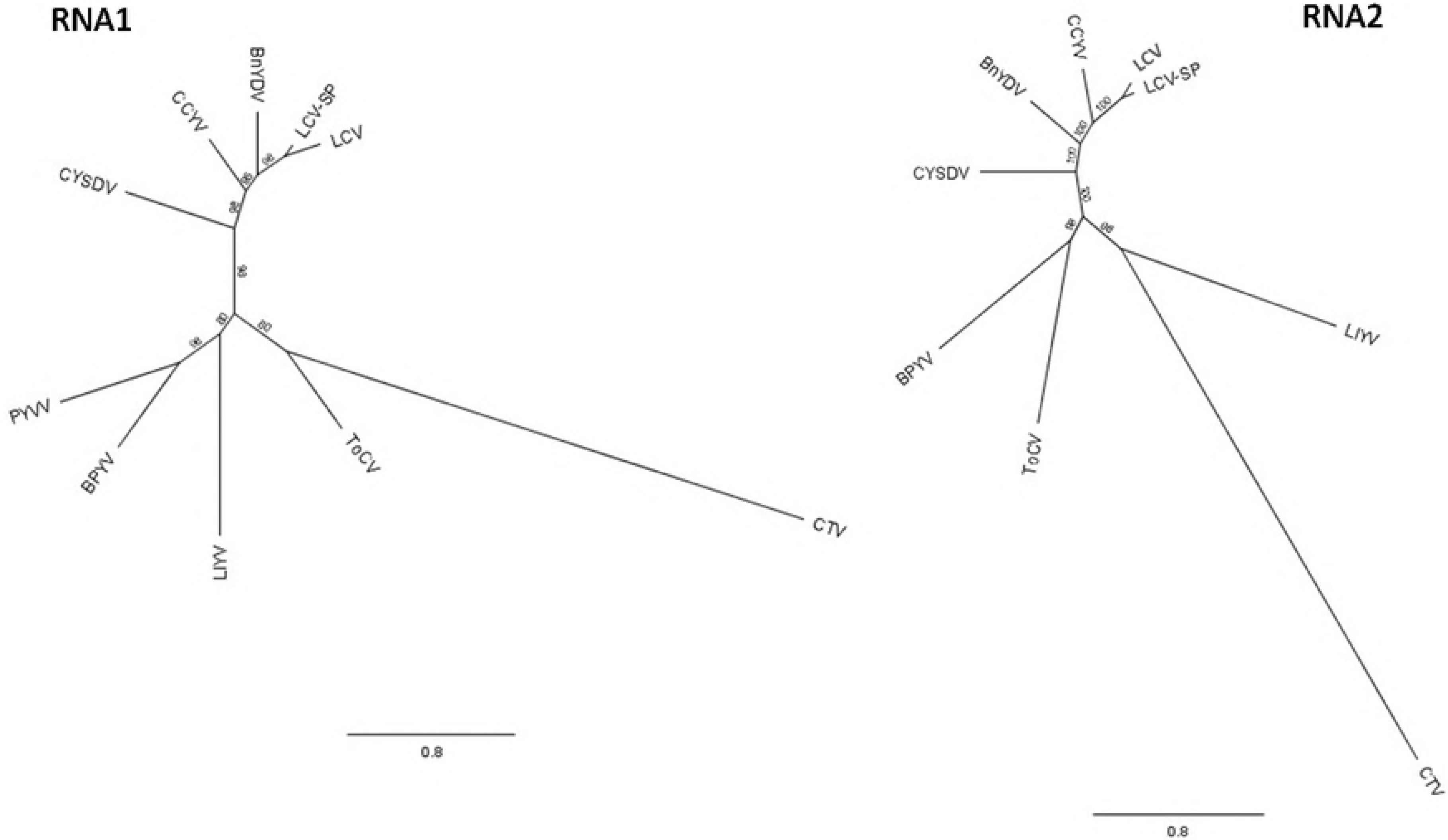
Bayesian phylogenetic trees of RNA1 and RNA2 full-length of selected viruses from the genus *Crinivirus*. *Citrus tristeza virus* (CTV, NC001661), was used as an outgroup. *Lettuce chlorosis virus* (LCV, FJ380118, FJ380119), *Lettuce chlorosis virus*-SP (LCV-SP, MG489894, MG489895), *Lettuce infectious yellows virus* (LIYV, NC003617, NC003618), *Cucurbit yellow disorder virus* (CYSDV, NC004809, NC004810), *Cucurbit chlorotic yellow virus* (CCYV, JQ904628, JQ904629), *Bean yellow disorder virus* (BnYDV), *Beet pseudo yellows virus* (BPYV, AY330918, AY330919), *Tomato chlorosis virus* (ToCV, KP137100, KP137101), *Potato yellow vein virus* (PYVV, NC006062). PYVV genome is tripartite and only its RNA1 has been considered in this analysis. Posterior probability is indicated.

Identical typology is observed when the Bayesian analysis is done for the RdRp region (Fig 4). When the analysis is done for other genomic relevant regions for *Crininvirus* species demarcation as CP and CPm, two clades are also generated where LCV-SP group with LCV, CCYV, BnYDV and CYSDV. Although in the same genetic clade, CYSDV is particularly distant from LCV-SP, LCV, CCYV and BnYDV. The topology generated in the phylogenetical analysis carried out for the HSP70 region was slightly different from the CP gene tree because LCV-SP was located nearer BnYDV than CCYV (Fig 4). This may indicate distinct recombination events in the *Crinivirus* genus. All the phylogenetic trees presented in this work show that, independently of the gene analysed, BnYDV and CCYV are closely related to LCV-SP and LCV although LCV, LCV-SP and CCYV, have different host ranges and also have not been described in the same geographic area.

**Fig 4.**
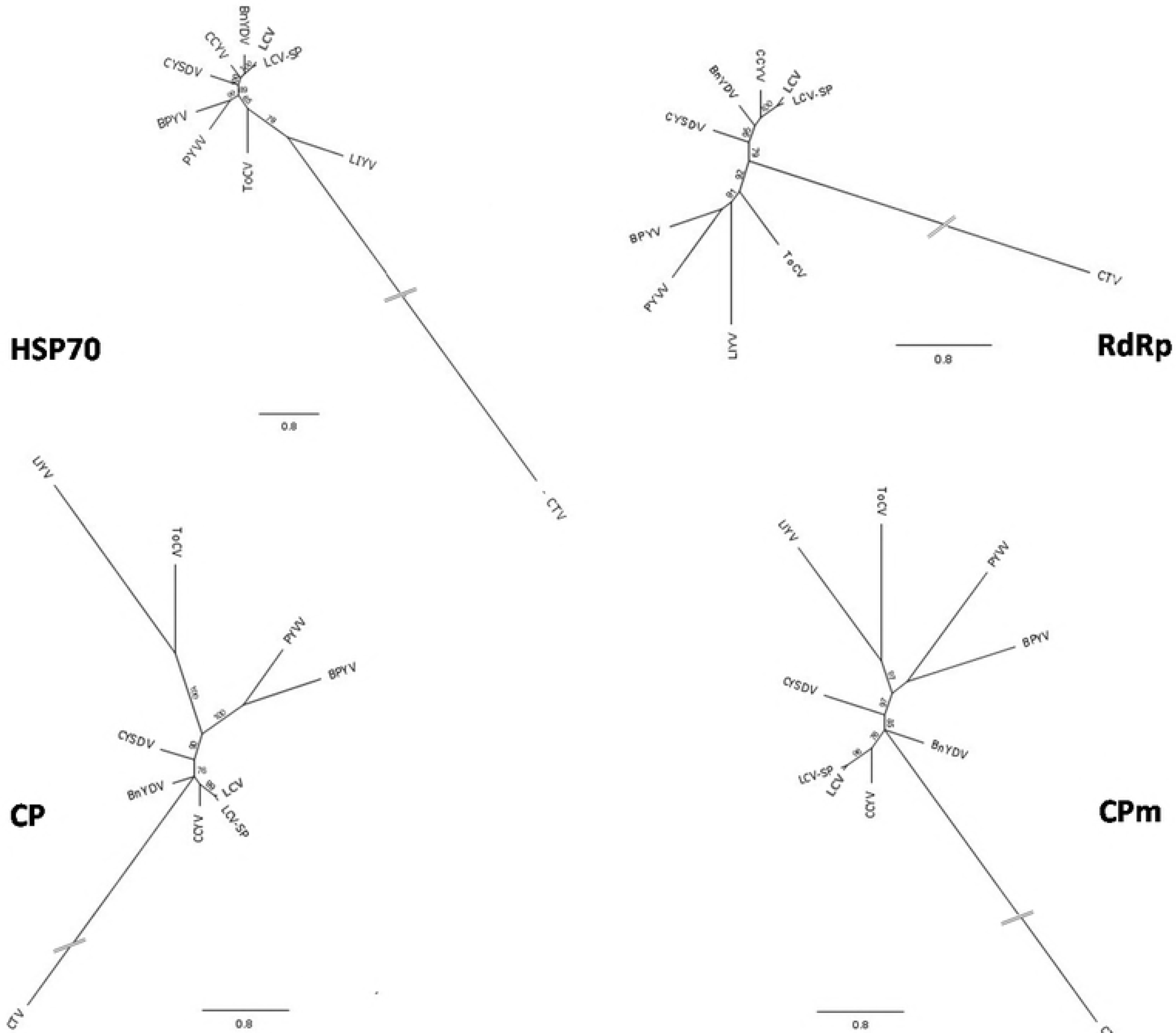
Bayesian phylogenetic trees for genomic relevant regions for *Crinivirus* genus demarcation as HSP70, RdRp, CP and CPm. Accesion numbers and virus acronyms are as in Fig 3. NC006063 and NC006061 corresponding with RNA2 and RNA3 of PYVV were added for HSP70, CP and CPm analysis.

### Recombination analysis and variability in the recombinant genomic region

RDP4 package program was used to further confirm the putative existence of first recombinant *Crinivirus* detected by Sanger and deep sequencing. Genomic sequence of RNA1 and RNA 2 belonging to LCV, BnYDV, CCYV and LCV-SP aligned were scanned in RDP4 using multiple methods. The full recombination scan of both RNA genomes (1 and 2) of LCV, BnYDV, CCYV and LCV-SP resulted in four recombination events detected (Figure 5, Table 2). The analysis detected LCV RNA1 and BnYDV RNA1 as possible major and minor parental sequences for LCV-SP RNA1 (event 1) with a 99% level of confidence (Table 2, event 1). This event was identified by all seven methods implemented in this package and, all of them show higher *p* value than any other algorithm executed in the rest of the events detected. BnYDV RNA1 was also suggested as a major parental sequence for CCYV RNA1 using five algorithms and with unknown minor parental (Table 2, event 2). Event 3 was detected by seven algorithms and signalized to LCV RNA2 as minor parental and CCYV-RNA2 as a possible recombinant. Although this event was suggested for seven methods as event number 1 was, the highest *p*-value (7,181 × 10−^9^, RDP) is much lower than in event 1 (5,710×10−^103^, RDP). LCV RNA2 was identified for six methods as a putative recombinant in event 4 with LCV-SP RNA2 and CCYV RNA2 as major and minor possible parental respectively (Table 2, event 4).

**Fig 5.**
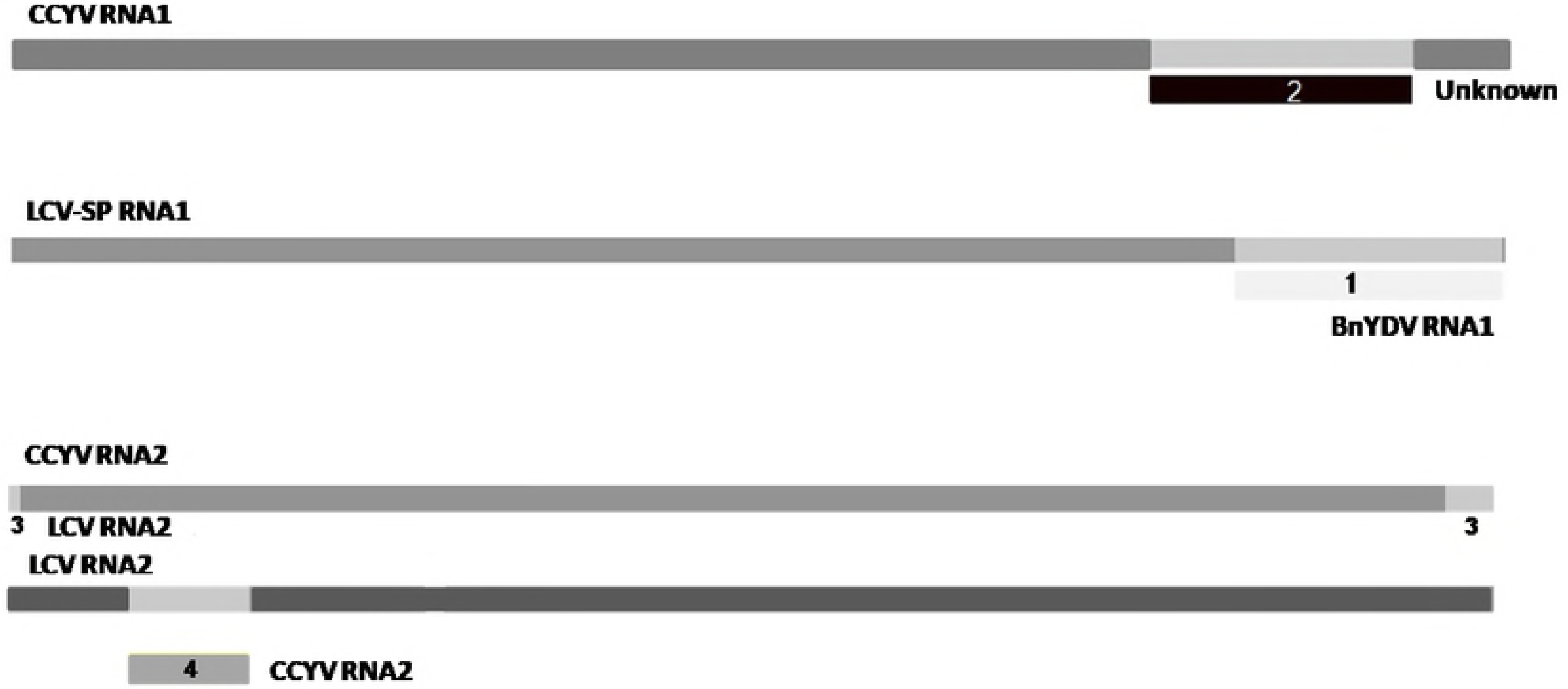
Recombination analysis of RNA1 and RNA2 of LCV, BnYDV, LCV-SP and CCYV using the recombination detection program (RDP-4).

**Table 2.** Unique recombination events identified by Recombination Detection Program v 4.80 (RDP4).

| Event | Recombinant | Major parent | Minor parent | RDP | Geneconv | Bootscan | Maxchi | Chimaera | Siscan | 3Seq | Begin | End |
| --- | --- | --- | --- | --- | --- | --- | --- | --- | --- | --- | --- | --- |
| 1 | LCV-SP RNA1 | LCV RNA1 | BnYDV RNA1 | 5,710 x 10<sup>-103</sup> | 6,418 x 10<sup>-63</sup> | 2,665 x 10<sup>-107</sup> | 5,521 x 10<sup>-29</sup> | 4,462 x10<sup>-28</sup> | 2,232 x 10<sup>-36</sup> | 1,332x10<sup>-15</sup> | 7580 | 8815 |
| 2 | CCYV RNA1 | BnYDV RNA1 | Unknown | 2,182 x 10<sup>-4</sup> |  |  | 1,403 x 10<sup>-2</sup> | 6,949 x 10<sup>-4</sup> | 3,625 x 10<sup>-5</sup> | 2,131 x 10<sup>-2</sup> | 7044 | 8225 |
| 3 | CCYV RNA2 | Unknown | LCV RNA2 | 1,498 x 10<sup>-9</sup> | 1,498 x 10<sup>-4</sup> | 3,316 x 10<sup>-9</sup> | 2,801 x 10<sup>-2</sup> | 4,708 x 10<sup>-4</sup> | 6,834 x 10<sup>-6</sup> | 7,181 x 10<sup>-9</sup> | 7858 | 75 |
| 4 | LCV RNA2 | LCV-SP RNA2 | CCYV RNA2 | 6,581 x 10<sup>-6</sup> | 2,614 x 10<sup>-2</sup> | 6,247 x10<sup>-3</sup> | 5,859 x 10<sup>-9</sup> | 5,244 x 10<sup>-6</sup> |  | 1,109 x 10<sup>-4</sup> | 585 | 1046 |

To confirm the result obtained from RDP4, BootScan analysis using each putative recombinant as query sequence were executed with the Simplot package. When the analysis was performed for LCV-SP RNA1, a point of recombination was detected in the same genomic area described by RDP4 (Fig 6). This result supports the evidence that LCV RNA1 (FJ380118) and BnYDV RNA1 (EU191904) are the origin of the LCV-SP RNA1 genome. The recombination events 2 and 4 however, were not recognized by the Simplot package. Bootscan analysis using CCYV RNA2 as query (Fig 7) suggested a recombination point at 3´UTR as is described in RDP4 (Fig 5) but not in the 5´UTR region (Fig 6).

**Fig 6.**
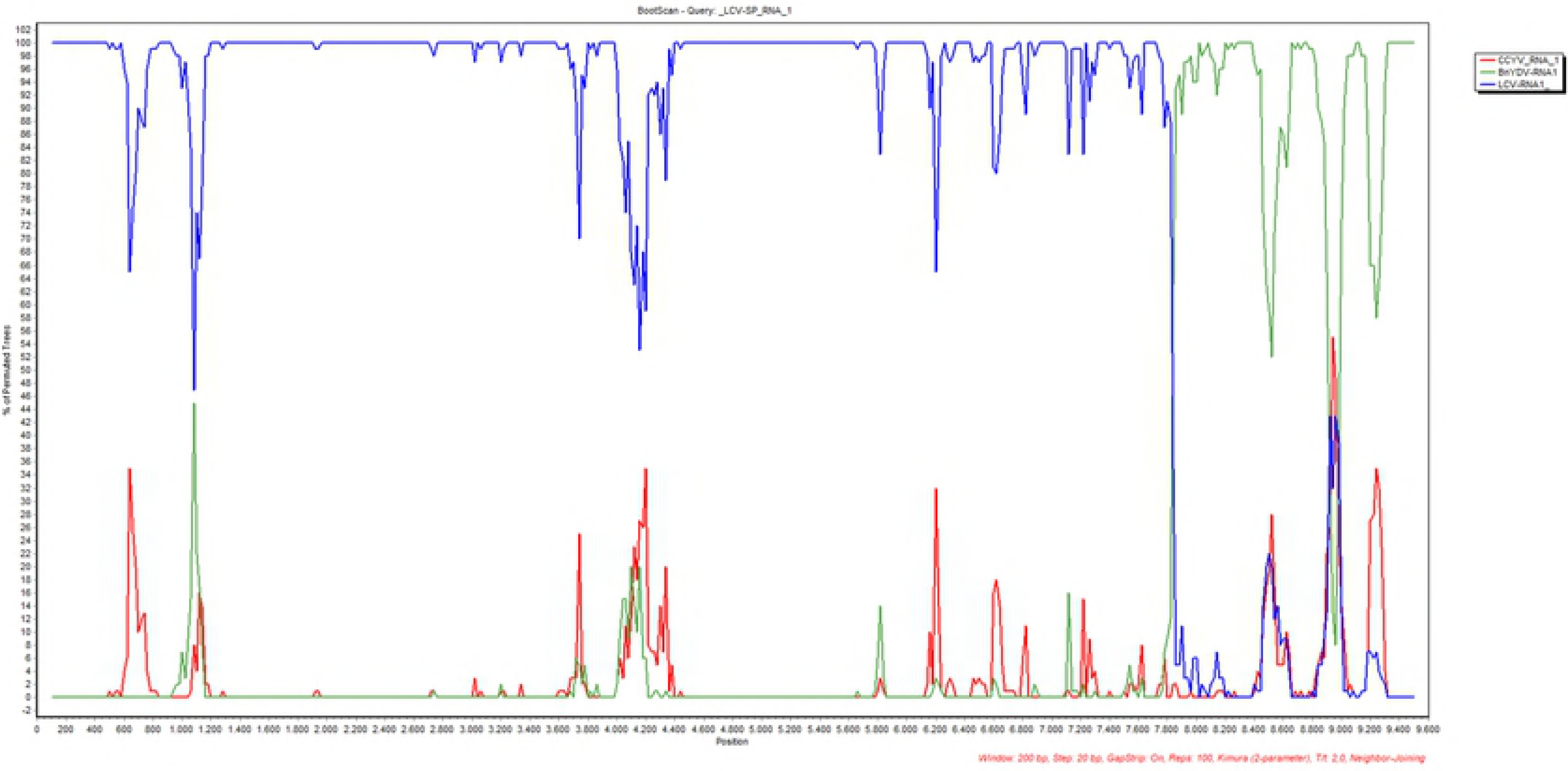
Bootscan analysis with Simplot program using LCV-SP RNA1 as the query sequence. CCYV, LCV, and BnYDV RNA1 were compared. Analysis was carried out with a sliding window of 200 bp and a step size of 20 bp.

**Fig 7.**
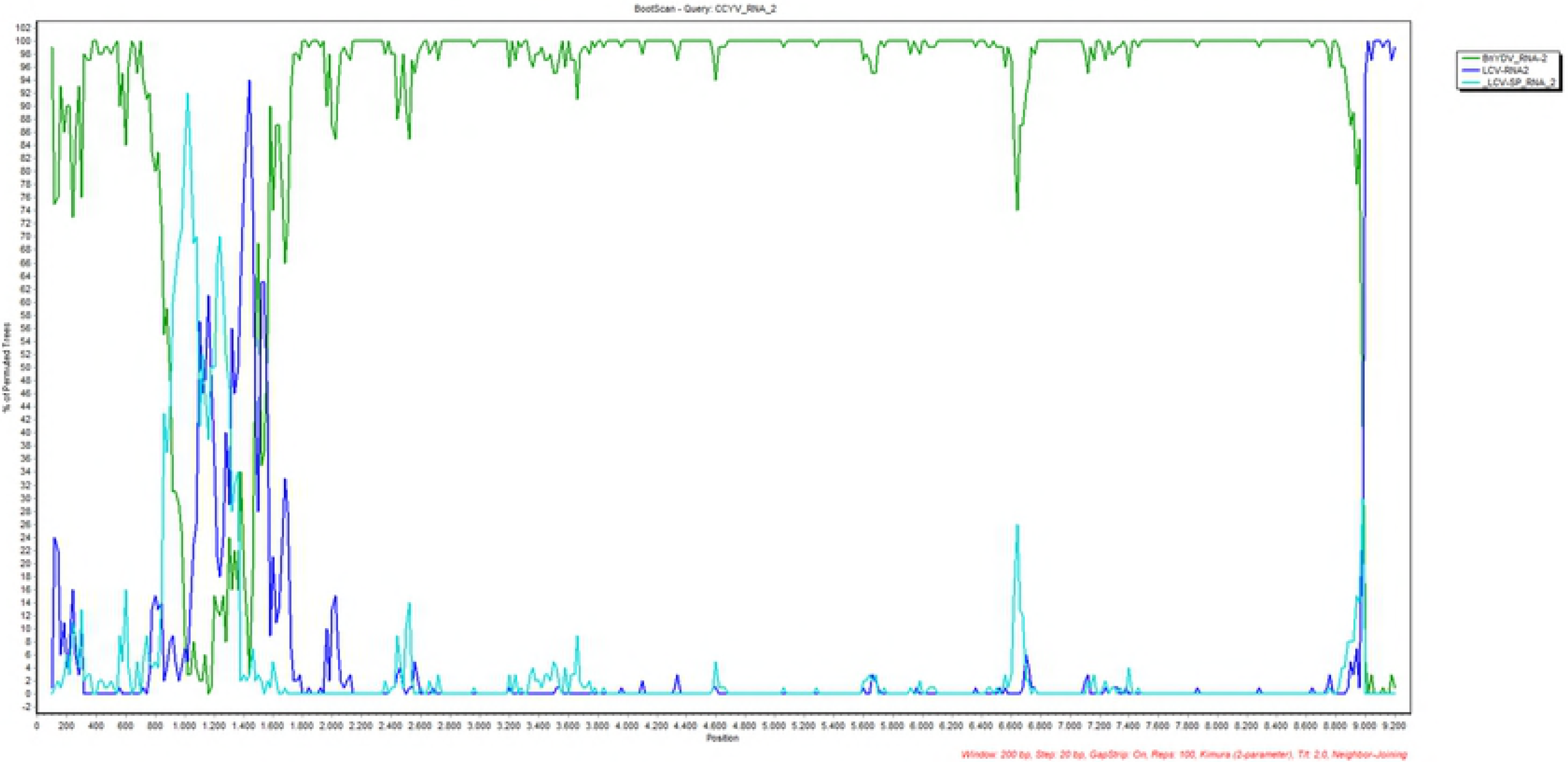
Bootscan analysis with Simplot program using CCYV RNA2 as the query sequence. CCYV, LCV, and BnYDV RNA2 were compared. Analysis was carried out with a sliding window of 200 bp and a step size of 20 bp.

Eleven DNA fragments corresponding with the entire P26 and P6 sequence of the RNA1 of LCV-SP were submitted to the GenBank database with the accession numbers assigned (MH170031-MH170041). These sequences confirmed the presence of the recombinant LCV-SP in the south of Spain since at least 2011. Values of genetic distance sequences at non-synonymous positions for both ORFs were of the same magnitude (Table 3). The ratio between nucleotide diversity values in non-synonymous and synonymous positions (dNS/dS ratio) indicates the amount of variation in the 18nucleic acid which results in variation in the encoded protein [34–36]. For P26, genetic distance at synonymous positions was null, and consequently, it was not possible to calculate the dNS/dS ratio. In the case of P6, The dNS/dS ratio was below unity, whic could indicate a negative selection against protein change. However, the value of the xstandard error is too high to consider the data conclusively.

**Table 3.**
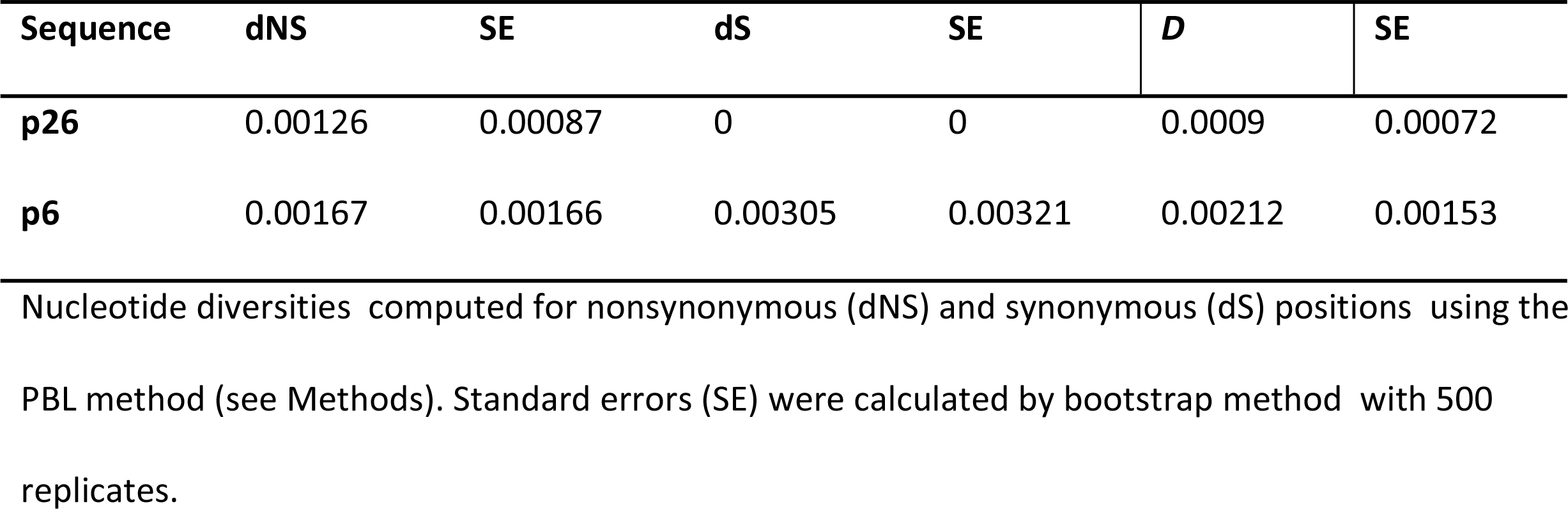
Nucleotide diversity for the 3´end LCVSP genome.

### Molecular differentiation between yellowing induced by BnYDV or LCV-SP

Restriction analysis with *Kpn*I after RT-PCR with the primers LC-BnF/LC-BnL, which amplified partial sequence of RdRp and P26 in both viruses, yielded two fragments of 582 y 157 bp when the virus was LCV-SP and a single band of 769 bp when the virus present was BnYDV. This RFLP test is then capable of distinguishing yellowing disease originating from recombinant or non-recombinant virus isolates (Fig 8).

**Fig 8.**
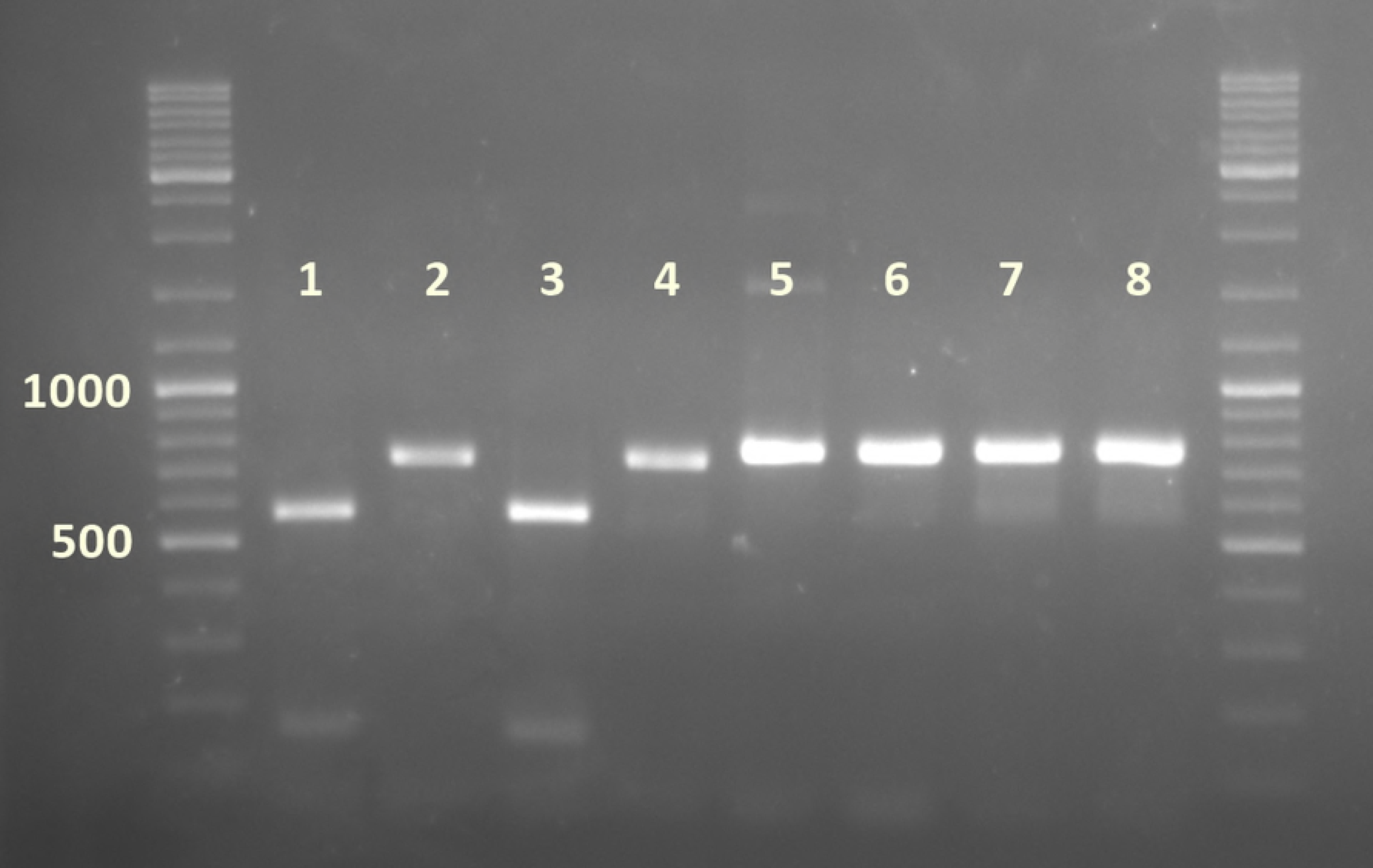
RFLP analysis with *Kpn*I after RT-PCR with by primers LC-BnF/LC-BnL. Lanes 1, 3: cut amplification products of LCV-SP isolates obtained in 2013 and 2017 respectively. Lanes 2 and 4: same amplification products uncut. Lane 5 and 7: cut amplification product of BnYDV isolates obtained in 2005 and 2009. Lanes 6 and 8: same amplification products uncut.

### Discussion

Members of genus *Crinivirus* represent a worldwide emerging disease as evidence of the increasing number of new species identified during the past 20 years [37–39, 9, 4]. Or, as in another example, a species already described increases its host range [40]. New LCV isolates have been recently described affecting ornamental plants, tobacco and tomato crops [41–43]; but there is no information about the capability of these LCV isolates to infect lettuce crops. Primer walking method and deep sequencing of vsRNA allowed the elucidation of the sequence and genome organization of the first recombinant closterovirus resulting after crossover recombination of intact ORFs. The 5´ from LCV-SP RNA1 includes the replication module (ORF 1a and 1b), which share high homology (more than 90%) with the LCV type isolate (FJ380118); however, the 3´end, which encompass P26 and P6, shares high amino acid identity with the 3´end of BnYDV RNA1 (99 and 100%, respectively). These ORFs, are described to be expressed in LCV from 3´ coterminal subgenomic RNAs (sgRNAs), a typical strategy among closteroviruses [8,6,44] that increases the probability of recombination events [45].

The genomic sequence of LCV-SP RNA2 share 8 ORFs with the Californian isolate of LCV (FJ380119); P6, and P4.8, located in the 5´and 3´end of the Californian isolate, respectively, and with unknown function, are absent in LCV-SP. Both putative proteins are also absent in the LCV Chinese isolates recently reported [41,42], which could indicate that these genes are not essential in the biology of LCV.

3´UTRs sequences in both genomic RNAs are highly conserved, sharing 72% of nucleotide sequence identity. This feature is common among criniviruses [46] and it agrees with the notion that sequence identity is essential to generate structures that facilitate the synthesis of the negative sense RNA during replication [47,48,44].

The accumulation of vsiRNAs observed in different genome regions probably involve gene expression strategies common within closteroviruses as translational frameshift and the production of 3´coterminal sgRNAs [8]. The lower presence of vsiRNAs along ORF 1b, compared with ORF 1a, suggest that this ORF is presumably expressed by translation via a +1 ribosomal frameshift rather than by sgRNA production as it was proposed before for LCV and other closteroviruses [49–51,6]. The presence of vsiRNAs hot spots along ORF 2, 3 and 4 of LCV-SP RNA2, could correspond with the production of sgRNAs, as was described in LCV [6].

Higher number of 21-nt class vsiRNAs populations with respect to 22-nt support the prevalence of a Dicer like protein 4 (DCL4) in the production of vsiRNAs in beans, as has been suggested in others plants [52,53]. 21− and 22-nt vsiRNAs populations, showed the same preference for sense or antisense polarity in RNA1 and RNA2, indicating that the mechanisms responsible for strand polarity are not dependent on the preference of DCL enzymes but on other factors specific to this virus species. The nucleotides present at the 5´ and 3´ terminal positions were investigated and resulted in an overall preference for A/U nucleotides. This prevalence is widespread among plant virus-pathosystems [54] and indicates the involvement of similar AGO complexes in the silencing of LCV-SP by green beans. Although the function of P26 and P6 in LCV-SP RNA1 (and hence BnYDV-RNA1) is unknown, the reduction of vsiRNA populations in this region could suggest that P26 or P6 interfered with the accumulation of vsiRNA. Recently, P23 of LCV, the closest homologous protein of P26, and also located at the 3´end of RNA1, has been described as a viral suppressor of RNA silencing (VSRs) and whose suppressor activity produces a reduction of siRNA [7]. To elucidate this hypothesis additional experiments are required.

Phylogenetic analysis grouped LCV-SP closely in the same lineage as LCV, CCYV and BnYDV as was already reported [9,6]. Recombination analysis completed with RDP4 and Simplot programs, demonstrates the existence of natural crossover recombination of intact ORFs between LCV-RNA1 and BnYDV-RNA1 (event 1). The recombinant virus, LCV-SP, shifted its host range by affecting green bean crops and not lettuce, its original host [11,5]. No other evident recombination events were detected in our analyses. Events 2, 3 and 4 have been discarded as possible recombinants although, alternatively they could be indicative of the close genetic relationship between LCV, LCV-SP, BnYDV and CCYV, as has been observed in the phylogenetic analysis. Plant virus with segmented genomes seem to frequently accumulate recombinants [45,55]. Within genus *Crinivirus*, which shows the lowest variability within the family *Closteroviridae* [19], only *Sweet potato chlorotic stunt virus* (SPCV) and *Beet pseudo yellows virus* (BPYV) have been reported as examples of recombination-mediated gene gain [39,21]. The existence of LCV-SP changes the perspectives of the possible epidemiological scenarios within the genus *Crinivirus*, an emergent genus and, as we have demonstrated, with more recombination possibilities than previously shown.

## Acknowledgments

The authors want to thank to Antonia Belmonte for her excellent technical assistance
and Vince Reid for his help in the grammatical revision of the manuscript.

## Author Contributions

Conceived and designed the experiments: LR, DJ. Performed the experiments: LR, AS, CG, LV Analyzed the data: LR, LV, DJ. Wrote the paper: LR, DJ.

## Supporting information

**S1 Table. Primers list used to confirm the LCV-SP genome by RT-PCR.**

**S2 Table. Numbers of reads homologies identified by BLASTN from contigs obtained from small RNAs and dsRNAs libraries.**

**S1 Fig. Percentages of nucleotide species at the 5´ and 3´ ends of redundant vsiRNAs derived from LCV-SP RNA1 (A) and RNA2 (B) according to the size of the reads.**

